# Effect of Weather and Soil Management on Medicinal Compound Concentration in East African Arid Acacias: A Simulation Study

**DOI:** 10.1101/2025.01.10.632480

**Authors:** Esther Wairimu Kinyua, Stephan Pietsch, Esther Muli, Vyacheslav Kungurtsev, Sospeter Neru, Kennedy Mugo

## Abstract

In African countries various tree spcies are a key part in Non Timber Forest Production (NTFP) economy, be it for fruit, resin collection or medicinal use. Among these species Acacias play an important role in the arid and semi-arid dry-lands over Africa. In this study we explore the potential of BioGeoChemical Modelling techniques to assess growth peformance of Acacia ocurrences within their natural environment, where they only may persist against rain-fed grassland vegetation when having access to seasonally changing groundwater tables. Within this study we assess the applicability of BGC-modeling techonologies to pastoral and AgroForestry settings within a semi-arid region in Kenya. Results show that effects of (i) changing hydrological patterns, (ii) fire and grazing regimes and (iii) plant use for resin production may be captured.

## 2 Introduction

Medicinal plants have been an integral part of traditional healthcare systems for centuries, particularly in African countries like Kenya, where indigenous knowledge of plant use is deeply rooted in local communities [40]. Moreover, these plants are not only valued for their medicinal properties but also play a vital role in biodiversity, ecology, and cultural heritage [41]. In Kenya, our country of focus in this paper, indigenous medicinal trees such as Prunus africana, Warburgia ugandensis, Croton megalocarpus, and Moringa stenopetala have long been utilized in traditional medicine for treating a wide range of ailments [27]. However, over-exploitation, habitat loss, and climate change pose significant threats to these valuable species[1]. By presenting research into the metabolic mechanisms responsible for medicinal compound production in these flora, we hope to promote programs that seek to protect and incentivize greater growing initiatives, realizing their economic and social welfare potential.

Among these medicinal plants, the Acacia family, such as the Acacia nilotica and Acacia senegal species, stand out due to their dual role as medicinal resources and ecological stabilizers, particularly in arid and semi-arid regions [10]. Acacia is known for producing bioactive compounds with anti-inflammatory, antibacterial, and antioxidant properties, which are widely used in traditional medicine [22]. At the same time, these trees contribute to environmental sustainability by improving soil fertility through nitrogen fixation, preventing soil erosion, and providing shade and shelter for various species [70].

Acacias are of the greatest importance for the dry lands of Kenya, which constitutes the vast majority, 90%, of the land in Kenya. These trees are the cornerstone of agriculture and livestock in the zones dubbed Arid and Semi-arid lands (ASAL). These drylands comprise a couple of Kenyan counties. They are found mainly in two ecological zones: one, Kenyan deserts/semideserts in Garissa, Wajir, Marsabit, Turkana; two, bushland and thickets in most of the drylands of Kenya. The latter are further divided into 3 (northern Acacia and Commiphora bushland and thicket, Southern Acacia and Commiphora bushland and thicket, Somali Acacia and Commiphora bushland and thicket). All these zones depend on the capacity of Acacias to enrich the mostly barren, degraded, and unproductive soils in these zones. Through the interaction of Acacia Rhizobia and Mycorrhizae with the soil, it receives nitrogen and other relatively immobile nutrients such as phosphoru [31][48][64]. In the Mimosodeae family, which is the bean / legume family, the Acacia tree is a protein powerhouse for livestock, wildlife, and humans. Acacia trees grow in tough climates and have shown to have some of the most powerful secondary plant metabolites on the planet - with ability to cure diseases.

The local communities, in arid areas, such as the Maasai, Kamba, and Samburu have used these plants as medicines for centuries, and they learned the ethnobotany from one generation to the next. Statistics shows that the most used trees medicinally in the dry-land are the trees in the Acacia family (mimosodeae/bean family) [66][2][45]

There is a wide array of Acacia species found in Africa, known for playing a key role in ecological succession by colonizing degraded lands [18]. These trees help restore soil fertility and can be integrated long-term into farming systems. While they offer numerous benefits, they are often unpopular due to their thorns and tendency to become invasive. Additional studies have shown that some Acacia species draw nitrogen from groundwater rather than the atmosphere, can produce more crude protein per hectare compared to grain crops, exhibit growth rings in their wood, and offer the potential for gum arabic to be a profitable cash crop [23] [70]. However, farmers are hesitant to plant Acacia trees unless the benefits can be financially rewarding. Given the more disperse ecological and social benefits of Acacia planting, an important goal for breeders is to develop Acacia varieties so that their direct economic appeal to farmers rises to become more commensurate to their aggregate social benefits, incentivize their planting. Apiculture in the acacia commiphora bushland and thicket is of high importance to the communities that keep bees to earn extra income. The favorite tree for the bee keepers is the Acacia mellifera. The diversity of bees is high, with over 50 species, in the acacia commiphora bushland and thicket zone[6][44][39].

Today the acacia is under attack and its popularity dwindling as demand for wood, charcoal/firewood (acacia being the favorite charcoal in Kenya), rises. Further, short-term agriculture prospects and rising towns like Ngong, Matuu, Kitengela, Nairobi, and Kiserian have led to the clearing of the acacia tree.

Popular dryland medicinal trees are also a favorite of livestock and wild animals like giraffe, gerenuk, kudu, dik-dik, elephant, and others. These valued acacia’s are Acacia gerrardii, Acacia senegal, Acacia nilotica, Acacia mellifera, Acacia xanthophloea, Acacia tortilis, Acacia elatior, Acacia/Faidherbia albinda, Acacia erioloba, Acacia polyacantha, Acacia hockii, Acacia lahai. They need scientific intervention to ensure their continued survival and survival of the drylands.

### 2.1 Contributions

In this paper, we select several specific Acacia plots for careful study. Collecting the necessary soil, plant measurements and climate data, we use state of the art software BGC-MAN to simulate the growth of the plant. We consider a number of scenarios, as far as both exogeneous changes in the aforementioned quantities as well as potential farmer inputs into the soil in order to properly understand their influence on both the overall growth of the plant as well as the production of medicinal compounds.

### 2.2 Modeling of Plant Metabolism - Background

#### Plant Metabolism Simulations

Plant metabolism simulations are an important tool to understand the complex interrelated biochemical processes that, under different environmental stresses, ultimately result in plant growth, development, and productivity. New approaches for computational modeling have offered firm grounds for simulating metabolic pathways in plants, thereby allowing research to investigate and predict metabolic functionality across tissues and conditions in plants. One important approach is the integration of multiscale metabolic modeling, combining detailed compartmentalized models at the organ scale with dynamic whole-plant models. This approach enables comprehensive analysis of carbon and nitrogen balance, thus underpinning metabolic and physiological processes from subcellular levels to overall plant growth and yield [24]. Such models have been applied to a wide range of plant species, including barley, and have proven their utility in agricultural research and crop improvement.

In addition, genome-scale metabolic modeling frameworks have recently arisen as some of the most powerful approaches to the analysis of whole plant systems. These frameworks are capable of simulating metabolic functions in multiple tissues, with a focus on optimizing resource utilization, such as photon capture for photosynthesis [14]. The incorporation of kinetic and regulatory factors into these models enhances their predictive capabilities, enabling more accurate simulations of plant responses to environmental changes. The development of synthetic metabolic pathways through computational approaches is another critical area of research. By employing optimization-based and retrobiosynthesis strategies, researchers can design and test synthetic pathways in silico, addressing gaps in the current understanding of plant metabolism [33]. This approach not only aids in metabolic engineering but also facilitates the exploration of novel biochemical pathways that could enhance crop resilience and productivity.

Furthermore, omics data integrated into metabolic models will be highly useful to enhance our knowledge regarding plant metabolic pathways. This can be achieved by employing mathematical and statistical models that integrate the genomic, transcriptomic, and proteomic levels of data to facilitate insights into the regulatory networks of metabolic processes [74]. It becomes of particular relevance in the development of strategies aimed at improving plant tolerance to abiotic stresses, such as drought and salinity, through the identification of key metabolic pathways and regulatory mechanisms involved in stress responses.

The development of plant metabolism simulations has been a rapidly evolving area, mainly due to rapid advances in computational modeling and integration of multi-omics data. These advances not only deepen our understanding of plant metabolic processes but also open new avenues for novel approaches toward crop improvement and sustainable agriculture.

Plant simulation models are important for understanding and predicting many aspects of plant growth and development. Models can be broadly divided into two categories: those that simulate the appearance and morphology of plants and those that focus on the growth processes and interactions with environmental factors. These models have mainly been influenced by the advancement in computational techniques and integration of biological data. The functional-structure modeling based on a fuzzy inference system is one important approach in plant simulation. Based on this approach, there is the ability to simulate plants’ appearance and predict growth status by considering the complex interactions between the plants and their environments, as proposed by Yin and Li (2014)[71]. The integration of geometric and physiological data into growth models has also enhanced the realism of simulations, enabling researchers to reproduce the growth processes of plants more accurately [35]. However, the complexity of plant structures and the need for extensive growth data pose significant challenges in model construction, often requiring sophisticated algorithms and computational resources.

Simulation models are used in industrial applications to optimize plant operations for better efficiency. In process engineering, tracking simulators calibrate plant states to streaming data in a dynamic fashion to First Principles Models (FPMs). Due to high cost of development and maintenance costs, more adaptive boostrapped dynamic models have been under development to achieve similar accuracy at less total cost [38]. In that respect, simpler simulation models can be used for real-time tracking even though they may not have the required accuracy for fine-tuning based on operational measurements. The complexity of the model and the difficulty of its use represent the limitation of this approach and a general problem in simulation modeling. Besides, the reliability of simulation models depends on proper calibration. According to literature findings, models calibrated with small sample size may not predict the outcome well; therefore, the uncertainties related to the modeling have to be considered carefully [8]. This is especially an issue in WWTPs, where the introduction of control components and actual behavior into simulation models does indeed improve their predictive power very significantly [36]. However, scarce data dependence remains a common problem for various types of plant simulations.

In the context of agricultural applications, simulation models like CropSyst have been developed to assess plant responses to varying environmental conditions and management practices [7]. These models facilitate the exploration of different crop management strategies, but they also face limitations in accurately capturing the complexities of plant-environment interactions. As such, ongoing research is essential to refine these models and improve their applicability to real-world scenarios.

Therefore, even though plant growth simulation models provide insights into both the process of growth and operational efficiencies, they also have a host of challenges. Some issues that need to be looked into include data scarcity, model complexity, and calibration to increase the dependability and applicability in agriculture as well as industry. It is expected that these will be overcome with future advances in computational techniques and integration of data.

Mathematical simulation models of plant metabolism are a very important tool for understanding the complex biochemical processes that govern plant growth, development, and responses to environmental changes. These models can be further divided into several types, each with its own methodology, application, and challenges inherent in their nature.

#### Types of Mathematical Simulation Models

1. Flux Balance Analysis (FBA): It is one of the widely used approaches that uses linear programming for the prediction of flux distribution in a network under a steady-state condition. The approach is highly appropriate in genome-scale metabolic models as it allows the modeling of different metabolic scenarios in order to optimize conditions of interest, such as biomass production or metabolite yield, as stated by Colombié et al., 2014, and Sweetlove & Ratcliffe, 2011[13][65]. FBA has been applied to the study of metabolic reprogramming in plants under different conditions, such as developmental stages and environmental conditions, using tomato fruit development [13] and Arabidopsis growth [62].
2. Dynamic Flux Balance Analysis – (dFBA): An extension of the previous work through the inclusion of time-sensitive aspects of metabolic flux and can be used to study dynamics like growth cycles or nutrient distribution in the whole lifespan of a plant, proposed by Shaw & Cheung, 2018, and Grafahrend-Belau et al., 2013 [62][24]. Indeed, this model proved successful in describing changes of metabolic states across growth phases under contrasting resource conditions.
3. Constraint-Based Modeling: This approach involves defining a set of constraints based on biological knowledge to predict feasible metabolic states. It has been utilized to analyze metabolic networks in various plant species, allowing for the integration of omics data to enhance model accuracy [60][19][9]. These models have been invaluable for understanding the interactions between plants and pathogens, as well as metabolic engineering applications [19][9].
4. Hierarchical and Multi-Organ Models: Hierarchical models, such as those based on k-core decomposition, provide insights into the organization of metabolic networks by analyzing the connectivity of metabolic reactions [21]. Multi-organ models simulate interactions between different plant tissues and organs, offering a comprehensive view of metabolic processes across the entire plant [24][46]. These models are crucial for studying systemic responses to environmental changes and genetic modifications.
5. Compartmentalized Models: This category of model allows the in silico simulation of metabolic compartmentalization, which is considered crucial for accurate modeling in plants due to their naturally compartmentalized cell environment, as stated by Pilalis et al. (2011) and Mintz-Oron et al. (2011)[55][42]. Such models allow researchers to consider a tissue-specific metabolic pathway and better understand the regulatory mechanism underlying plant metabolism in different organs.

#### Application of Mathematical Simulation Models

Plants’ metabolism mathematical models can be applied in the following areas:

- Metabolic Engineering: Models provide guidelines on the optimization of metabolic pathways for improving the yields of value-added compounds like pharmaceuticals and biofuels[63][33]. Genome-scale models, for example, have been used in engineering plants to enhance the production of a specific metabolite. [46].
- Breeding and Genetic Manipulation: By simulating metabolic responses to genetic modifications, researchers can identify potential targets for breeding programs aimed at improving crop yield and resilience [13][34].
- Understanding Plant-Environment Interactions: Models help in ascertaining how plants adapt their metabolism due to environmental stresses, including nutrient availability and climate change [69][25]. Such an understanding is important in developing strategies that will improve agricultural sustainability.

#### Challenges and Limitations

Despite the usefulness of these models, mathematical simulation models face some challenges that include:

- Complexity of Plant Metabolism: Due to the intricate nature of the metabolic networks, interactions, and feedback loops of plant metabolism, accurate modeling has often been challenging [25][75]. This is even more highly compartmentalized and therefore much more complex when it comes to the integration of metabolic pathways.
- Data Limitations: The accuracy of models heavily depends upon the availability and quality of experimental data. Knowledge gaps in metabolism and incompleteness of datasets are sources of inaccuracies in model predictions. Such has been reported by Baghalian et al. (2014)[9] and Hill et al. (2015)[25]. Further, integrating different omics data remains a big challenge.
- Validation and Predictive Power: While models can provide valuable insights, their predictive capabilities must be validated against experimental data. Discrepancies between model predictions and experimental results can undermine confidence in the model’s utility [42]. Continuous refinement and validation are necessary to improve model reliability.

Mathematical simulation models provide a powerful means to understand and optimize plant metabolism. With further increases in computing power and improvement of the experimental basis, such models will be increasingly more predictive and permit an effective, targeted management of crops, engineering of metabolic traits, and improvement of environmental resilience of crops. Realizing such potential, however, would presuppose the resolution of existing challenges related to model development, data availability, and their validation for basic and applied aspects of plant research.

## 3 Medicinal Acacias

Acacias have a long phylogenic history, and in fact there was a recent break of Acacias, in around 2012, those from Africa and those from Australia. Those in Australia, with no thorns, kept the name Acacia. And those in Africa, with thorns, were separated to two distinct categories: one, those having long thorns (Vachellia); and two, those with short thorns (Senegalia). For purposes of this study, we will stick to the name “Acacia” referring to all the African Acacias.

For this study, we have selected Acacia senegalensis, a species renowned for its production of valuable gum. It is native to areas such as Kajiado County, Kenya, and is well-suited to thrive in nutrient-deficient soils, where it plays a crucial role in enhancing soil fertility and supporting ecosystem stability.

### 3.1 Importance of Medicinal Trees in Kenya

Kenya is home to a diverse range of medicinal plants that form the backbone of healthcare for rural communities [47]. The reliance on these plants stems from their accessibility, affordability, and efficacy in treating common diseases such as malaria, skin infections, digestive disorders, and respiratory ailments [49]. For centuries, traditional healers have used indigenous knowledge to harness the medicinal properties of these trees, creating remedies that have been passed down through generations [27].

In addition to their medicinal value, these trees have economic significance, particularly in rural areas where commercialized herbal products are becoming an important source of income [12]. For example, the extraction and sale of medicinal bark, leaves, and gums from species like Prunus africana and Acacia senegal contribute to both the local economy and international trade [41].

### 3.2 Threats to Indigenous Medicinal Trees

Despite their importance, many medicinal trees in Kenya face critical threats [10]. Unsustainable harvesting practices, such as the over-collection of bark and roots, have led to population declines of key species. Deforestation and land conversion for agriculture further exacerbate the loss of natural habitats, leaving many medicinal tree species vulnerable[1]. Climate change also plays a role by altering the growth conditions and distribution of these trees, potentially reducing their medicinal potency and availability[73].

Acacia species, while relatively resilient, are not immune to these threats. Although they are adapted to thrive in arid and semi-arid environments, changes in rainfall patterns and increased demand for land have led to a decline in their populations [73]. This raises the need for strategies that ensure the sustainable management of Acacia and other medicinal trees while maintaining their medicinal value [1].

### 3.3 Focus on Acacia Species: Medicinal and Ecological Significance

Acacia species hold a unique position within Kenya’s medicinal flora due to their wide-ranging uses and ecological importance. Traditionally, Acacia nilotica has been used to treat conditions such as colds, coughs, and skin infections, while Acacia senegal is well known for its production of gum arabic, a substance used in pharmaceuticals, food, and cosmetics [70]. The medicinal compounds found in these species, such as tannins, flavonoids, and saponins, have been extensively studied for their therapeutic potential, particularly their anti-inflammatory and antimicrobial effects [22].

Beyond their medicinal value, Acacia trees play a critical role in maintaining the health of ecosystems, especially in semi-arid regions [16]. Fixing nitrogen in the soil, they improve soil fertility and contribute to agricultural productivity. Their deep root systems help prevent soil erosion, and their ability to survive in harsh climates makes them vital for restoring degraded landscapes [29].

Given these attributes, Acacia serves as an ideal model for studying the metabolism and growth of medicinal trees under various environmental conditions [12]. Understanding how Acacia responds to environmental stressors, such as drought and nutrient-poor soils, can inform the sustainable management of these species and ensure their continued use in traditional medicine [18].

Antiviral activity from in silico and molecular dynamics studies [20] Modeling of solar drying of gum in Acacia Senegal [5]

## 4 Previous Simulation Work of Medicinal Acacia Metabolism

Medicinal Acacias, have been studied comprehensively through various experimental and simulation approaches. Several studies highlight the impact of environmental factors and computational modeling on understanding Acacia metabolism. For instance, Mopipi et al. (2009) [43] investigated the effects of moisture, nitrogen availability, grass competition, and simulated browsing on the survival and growth of *Acacia karroo* seedlings. Their work gave emphasis to the interplay of abiotic and biotic factors in seedling establishment, which is crucial for modeling growth dynamics in varied environments. Ramoliya et al. (2004) [58] have also explored how soil salinization impacts the growth and nutrient accumulation of *Acacia catechu* seedlings. Their findings demonstrated significant reductions in uptake of macro- and micronutrient under saline conditions, which provides baseline data for simulating stress responses in Acacia metabolism. Traoré et al. (2007) [68] have also highlighted the role of Acacia species in enhancing soil carbon and nitrogen content in Sudanese soils. They connected these enhancements to increased soil respiration and microbial biomass, underlining Acacia’s ecological contributions and its carbon cycling role.

When it comes to simulations, Kirschbaum et al. (2008) [30] conducted simulations and scenario analyses to study soil carbon and nitrogen dynamics following the clearing of Mulga (*Acacia aneura*) vegetation in Queensland, Australia.

Their work has provided insights into post-disturbance nutrient fluxes and soil organic matter changes, which is critical for modeling long-term ecosystem responses. Bossel and Schäfer (1989) [11] also developed a generic simulation model for forest growth, which is comprised of carbon and nitrogen dynamics. Their application to tropical Acacias gives a framework for understanding plant-soil interactions and nutrient cycling across diverse ecological contexts. In addition, Sharpe et al. (1985) [61] employed a physiologically based continuous-time Markov model to simulate plant growth in semi-arid woodlands. This approach integrates stochasticity in environmental conditions, providing a robust methodology for studying Acacia growth in resource-limited settings. All of the above studies demonstrate the integration of experimental and simulation methodologies in advancing our understanding of Medicinal Acacia metabolism. The insights into environmental interactions, nutrient dynamics, and modeling frameworks collectively inform strategies for the sustainable management and conservation of Acacia species.

## 5 Details of the Current Study

### 5.1 Species Chosen for Current Study

Our species of choice *Acacia senegalensis*,is key in East Africa’s arid ecosystems. It is known for its ecological adaptability and valuable gum production. It is also found in regions of Kajiado County, Kenya. This species is able to thrive in nutrient-poor soils and contributes significantly to soil fertility and ecosystem stability. Its medicinal properties, influenced by weather conditions and soil management practices, accentuate the importance of understanding how these environmental factors affect the concentration of its medicinal compounds.

*Acacia senegalensis* plays a crucial role in East Africa’s arid ecosystems, where its ability to fix atmospheric nitrogen significantly enhances soil fertility. The litter falling from Acacia trees enriches the soil with organic matter, improving its nutrient content and moisture retention, which is vital in arid regions prone to nutrient depletion due to high evaporation and limited rainfall [4]. Under the canopy of Acacia, soil nitrogen and water content are higher than in open areas, supporting the growth of other plant species and contributing to overall biodiversity [4][26]. The interaction between Acacia roots and soil microbial communities further promotes ecological balance by fostering beneficial bacteria that enhance nutrient cycling, which is critical for both plant health and the biosynthesis of medicinal compounds [72][59][52].

Environmental factors, particularly rainfall and temperature, have a profound impact on the medicinal properties of *Acacia senegalensis*. Changes in precipitation can alter the physiological stress levels of plants, influencing the production of secondary metabolites. For example, drought conditions can lead to higher concentrations of certain medicinal compounds, while excessive rainfall can dilute or modify these compounds through changes in biosynthetic pathways [51][15]. Similarly, saline conditions, common in arid regions, can trigger adaptive mechanisms in the plant, which affect its growth and the concentration of bioactive compounds [3][57].

Soil management practices also play an essential role in optimizing the growth and medicinal potential of *Acacia senegalensis*. Conservation tillage, organic amendments and techniques such as mulching and Zai pits improve soil structure, improve water retention, and create a favorable environment for beneficial soil organisms, which in turn support plant health and synthesis of medicinal compounds [17][56][28]. The use of cover crops and integrated soil fertility management strategies further improve soil quality, ensuring a sustainable habitat for *Acacia senegalensis* in regions such as Kajiado County, Kenya [52][37][32].

Therefore, we can say that *Acacia senegalensis* is a vital species in arid ecosystems. Its ability to improve soil fertility, adapt to environmental stressors, and interact with microbial communities directly influences its medicinal value. Sustainable agricultural practices that improve soil health and manage water effectively are essential for maximizing the plant’s growth and bioactive compound production. Future research should investigate the interactions between soil quality, weather variability, and the biochemical pathways responsible for the synthesis of medicinal compounds in *Acacia senegalensis*, ensuring its continued ecological and medicinal significance.

### 5.2 Description of Site

We chose a typical AgroForestry environment close to Nairobi, i.e. the Kaijado site (1°31’S, 36°56’ E; 1580m a.s.l) (Fig. 1), that comprises typical ecotypes of semi-arid dry lands in landscapes of eastern, southern and western Africa. The site comprises of C3 and C4 grasses in savnnahs, and tree cover of *Acacia sengalensis* along hydrologically favorable subsites created by non-permanent brooks (Fig. 2). Dominant ecotypes of the site are mixtures of C3 and C4 grasses, accompanied by permanent tree cover along the non-permanent brook (cf. Fig. 2, A-C). The site is subject to extensive grazing and occasional fire events in the grasslands part, while permanent tree cover only occurs along the non-permanent brook, indicating water supply to the trees from the groundwater stream even during peak dry season.

**Figure 1:**
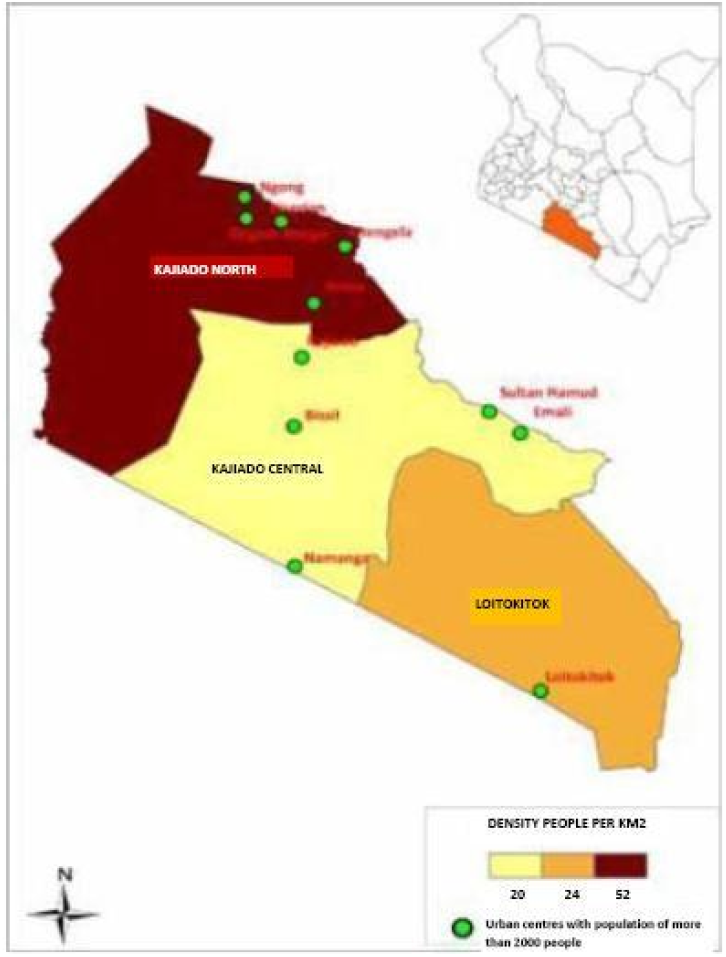
Map of Kajiado county[50]

**Figure 2:**
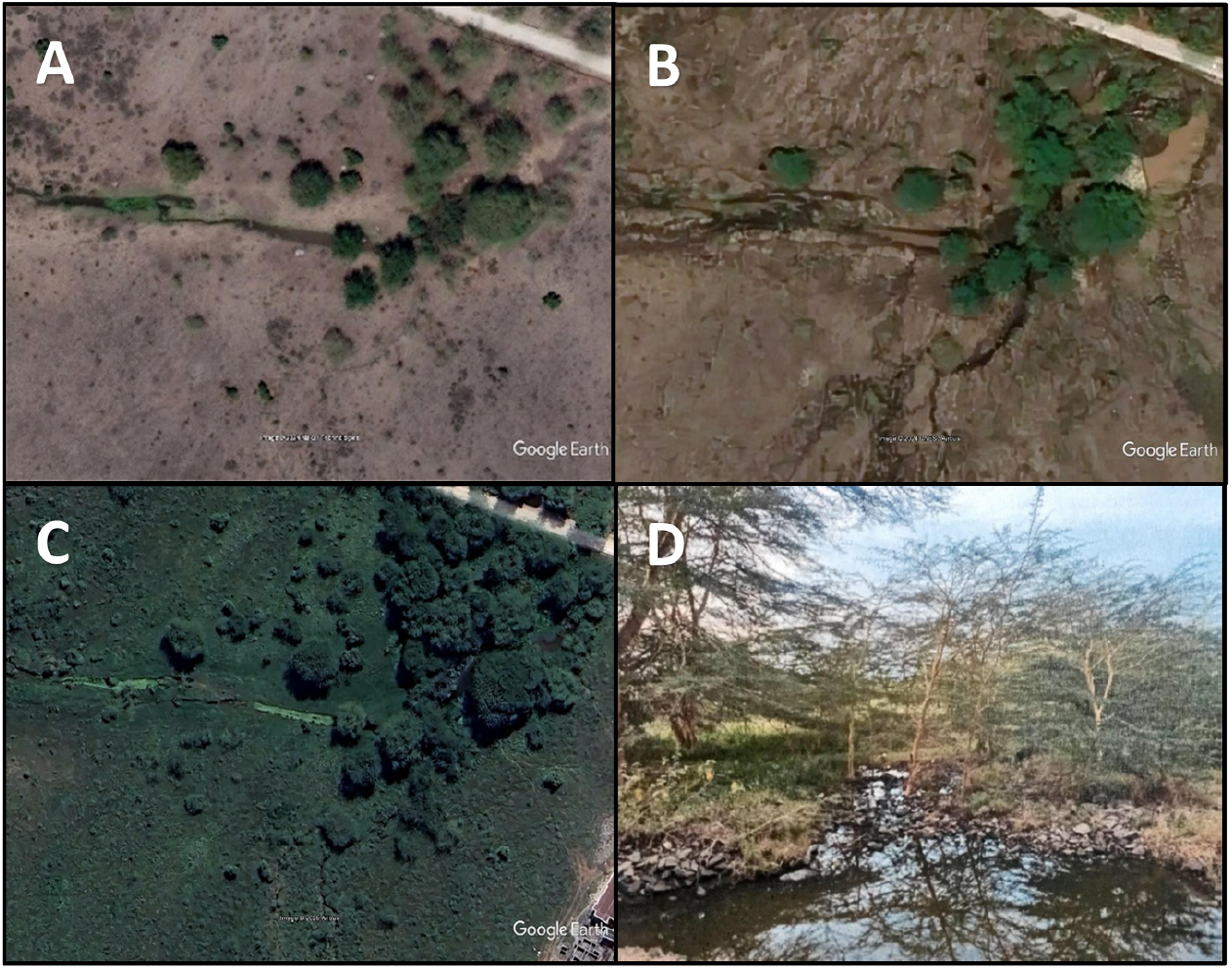
Different stages of the ecotypes present at the Kaijado site during different stages of seasonal development. A: 4 weeks after the onset of the dry season. B: end of the dry season. C: 4 weeks after onset of the rainy season. D: picture taken from the non-permanent brook towards the north-west at time of the development stage depicted in A

**Figure 3:**
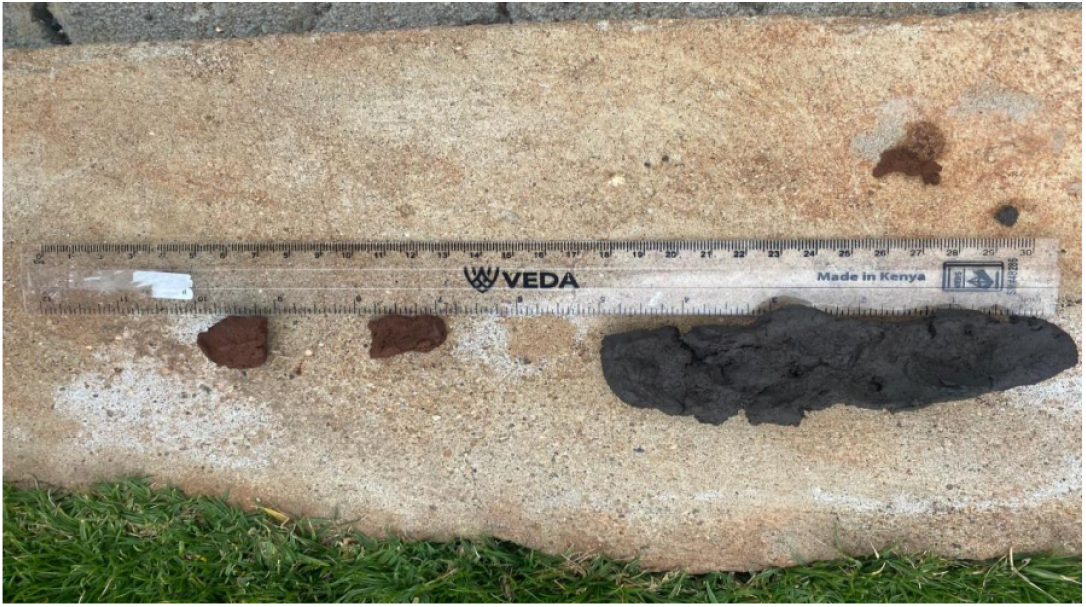
Ribbon Test Result: Sample on the right hand side (The soil ribbon was 5 inches/12cm making it clay soil. The soil very dark in colour almost black)

**Figure 4:**
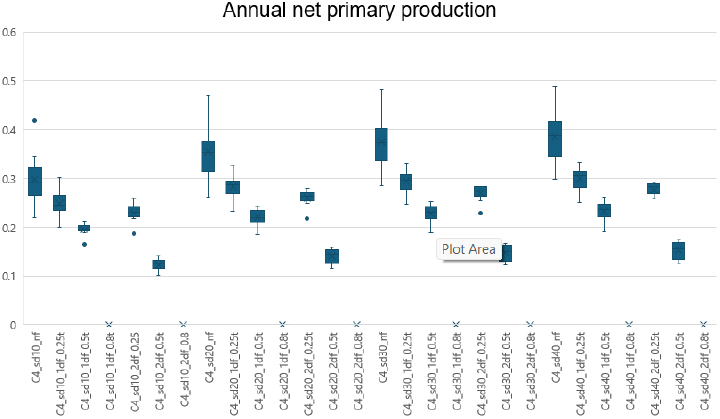
Enter Caption

**Figure 5:**
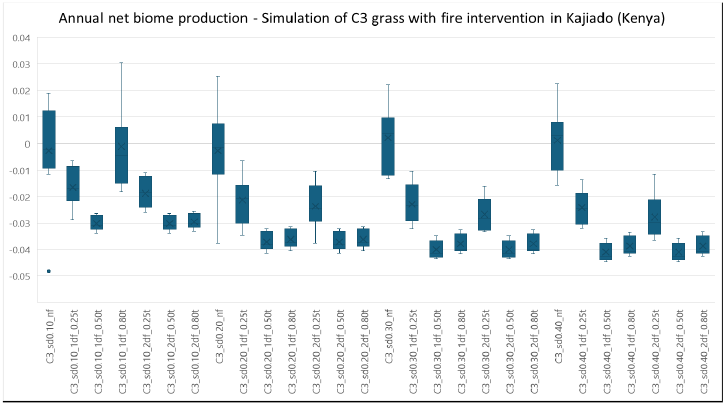
Enter Caption

**Figure 6:**
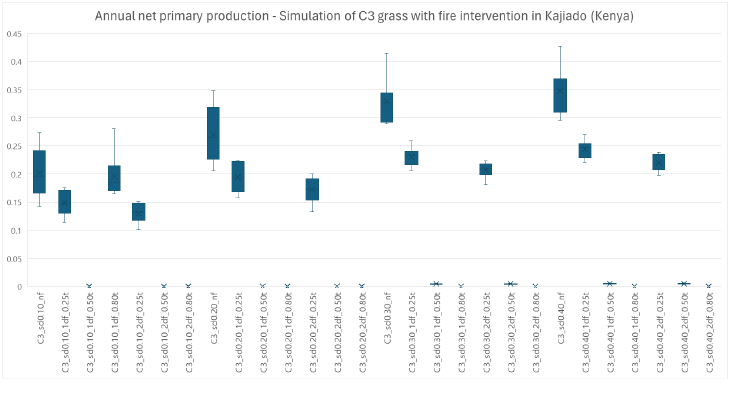
Enter Caption

**Figure 7:**
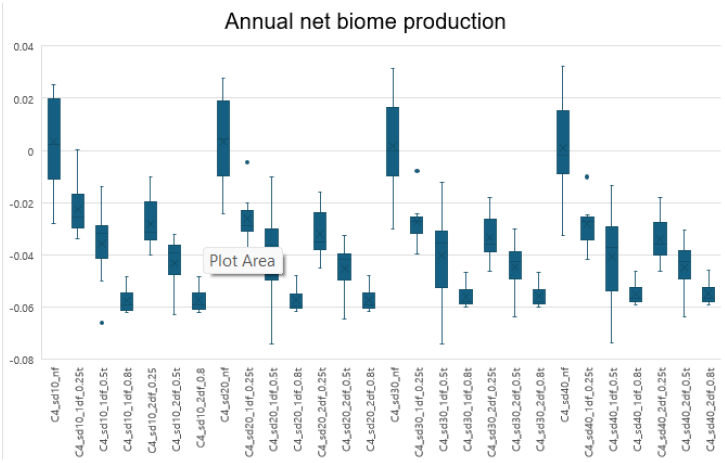
Enter Caption

**Figure 8:**
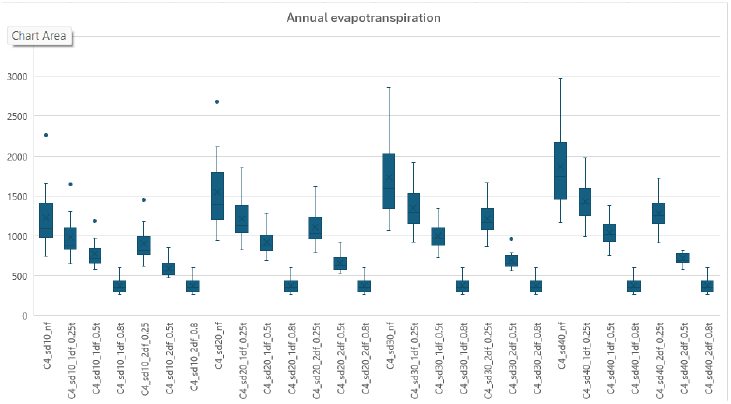
Enter Caption

**Figure 9:**
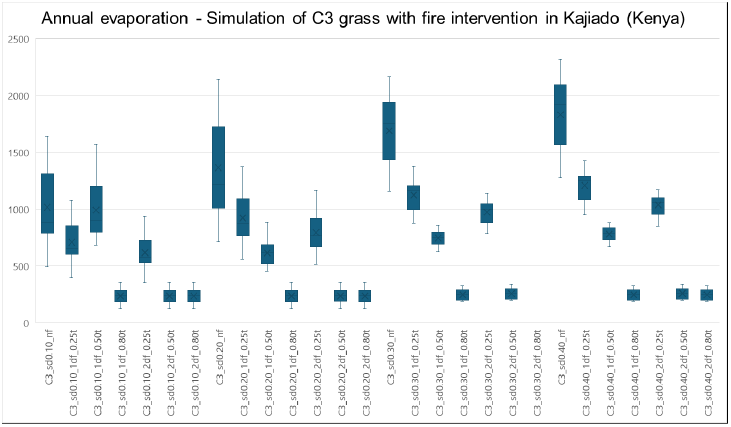
Enter Caption

### 5.3 Soil Characteristics

The study area exhibited a range of soil textures, each influencing the distribution and occurrence of different Acacia species. The soil textures were assessed using Finger and Ribbon tests, which provided understandings into the composition and suitability of soils for various species.

The soil in Kajiado County, particularly near the seasonal riverbanks, was predominantly black cotton clay, as indicated by soil tests (figure 2). The ribbon test produced a five-inch-long ribbon, and the finger test showed the soil to be sticky, buttery, and capable of molding like plasticine when wet. These characteristics confirm the presence of clay soil, known for its high water retention and poor drainage. The site was characterized by *Acacia senegalensis* thriving along the riverbanks, where moisture is relatively accessible, and *Acacia drepanolobium* found farther away, in drier areas.

Silt loam soils, which contain a higher proportion of clay, were also found in areas with slightly more clayey regions and lower elevations. These soils are particularly favorable for *Acacia senegalesnsis*, as they offer better moisture retention, which is crucial for survival in semi-arid environments. Silt loam, which also supports species like *Acacia mellifera* in mid-elevations with higher moisture retention, accentuates the ecological versatility of *Acacia senegalensis* in thriving under nutrient-poor, heavy clay soils while still contributing significantly to soil fertility and ecosystem resilience.

Sandy silt soils were also identified in parts of Kajiado County, which added to the region’s soil diversity. These soils are characterized by their moderate drainage and texture, which balances fine particles of silt and coarser sand.

Such conditions are beneficial for plants like *Acacia senegalensis*, as they provide better aeration and water retention compared to pure sand, while avoiding the waterlogging issues associated with heavy clay. This soil type further supports the adaptability and growth of *Acacia senegalensis*, especially in semi-arid environments like Kajiado.

#### Implications for Species Distribution

The variety in soil textures across Kajiado region plays a significant role in shaping the distribution patterns of Acacia species. *Acacia senegalensis* has shown to thrive in silt loam soils that offer higher moisture retention. The specific soil requirements and preferences of each species therefore are crucial in determining their occurrence in relation to elevation, highlighting the importance of soil characteristics in the ecology of the region.

## 6 Simulation - Background

### 6.1 Software Description

For this study, we used `BioGeoChemistry Management` (BGC-MAN) (Pietsch 2013) to assess the growth dynamics of acacia species and surrounding ecosystems, as they may depend on climate, soil hydrology and management. Originally begun as an earlier variant at Oak Ridge National Labs in the USA, it represents one of the novel groundbreaking Next Generation Vegetation Models, adaptable software that permits flexible parameter choices. Daily weather data and site information are used to calculate carbon, nitrogen, water, and energy cycles, allowing simulations to estimate the impact of possible land use management interventions.

Leaf area index (LAI, m2 leaf area/m2 ground area) is calculated by multiplying carbon allocated to leaves S.A. Pietsch et al. / Forest Ecology and Management 211 (2005) 264–295 265 times the specific leaf area (m2 leaf area/kg leaf carbon) and controls canopy radiation absorption, water interception, photosynthesis, and litter inputs to detrital pools. Net primary production (NPP) is based on gross primary production (GPP) calculated with the Farquhar photosynthesis routine (Farquhar et al., 1980) minus the autotrophic respiration. The autotrophic respiration includes the maintenance respiration, calculated as a function of tissue nitrogen concentration (Ryan, 1991), and growth respiration which is a function of the amount of carbon allocated to the different plant compartments (leaf, root and stem). NPP is partitioned into the leaves, roots and stems as a function of dynamic allocation patterns, considering possible limitations in the availability of nitrogen.

The model requires meteorological input data, such as daily minimum and maximum temperature, incident solar radiation, vapor pressure deficit (VPD), and precipitation. Aspect, elevation, nitrogen deposition and fixation, and physical soil properties are needed to calculate: daily canopy interception, evaporation and transpiration; soil evaporation, out-flow, water potential and water content; LAI; stomatal conductance and assimilation of sunlight and shaded canopy fractions; growth and maintenance respiration; GPP and NPP; allocation; litter-fall and decomposition; mineralization, denitrification, leaching and volatile nitrogen losses.

Besides changes in site and climate conditions, BGC-MAN explicitly considers ecosystem interventions like seasonal changes in soil hydrology ([54]), the effects of harvesting of ecosystem products by animal grazing or direct human intervention ([67], or any combinations of them.

### 6.2 Simulation Procedure

We followed a standard simulation protocol ([54] starting from a dynamic steady state ([53]) for each ecotype, followed by current land use practices from 1950 to 2023. For the savnnah ecotypes of the Kajiado site we simulated rain-fed growth performance in dependence on effective soil depth and impacts of fire occurrence(1-2 days) and intensity (0.25, 0.5, 0.8 fraction of biomass turnover). For tree cover along the nonpermanent brook, we compared rain-fed growth performance with two different ground-water access scenarios (July-December, August to November).

resin collection scenarios

The assessment of the impact of resin collection included stem tissue harvesting events, followed by changes in plant allocation patterns in reaction to changed metabolic demands (Petritsch et al. 2008?). Specifically we changed the ratios of above to belowground and stem to leaeves allocation patterns as follows …

## 7 Simulation Results and Observations

Grassland productive depends largely on seasonal rainfall and is affected by grazing and fire, acacia productivity and growth exhibited a pronounced dependence on soil water hydrology.

resin collection - a few more scenarios 2-4 of 2-4 split graphs

## 8 Conclusion

**REFERENCES ON INDIGENOUS MEDICINAL TREES STUDY WITH A FOCUS ON ACACIA**

